# NINJA-Associated ERF19 Negatively Regulates *Arabidopsis* Pattern-Triggered Immunity

**DOI:** 10.1101/180059

**Authors:** Pin-Yao Huang, Jingsong Zhang, Beier Jiang, Jhong-He Yu, Yu-Ping Lu, KwiMi Chung, Laurent Zimmerli

## Abstract

Recognition of microbe-associated molecular patterns (MAMPs) derived from invading pathogens by plant pattern recognition receptors (PRRs) initiates defense responses known as pattern-triggered immunity (PTI). Transcription factors (TFs) orchestrate the onset of PTI through complex signaling networks. Here, we characterize the function of ERF19, a member of the *Arabidopsis thaliana* ethylene response factor (ERF) family. ERF19 was found to act as a negative regulator of PTI against *Botrytis cinerea* and *Pseudomonas syringae* pv. *tomato* DC3000 (*Pst*). Notably, overexpression of *ERF19* increased plant susceptibility to these pathogens and repressed MAMP-induced PTI outputs. In contrast, expression of the chimeric dominant repressor *ERF19-SRDX* boosted PTI activation, conferred increased resistance to *B. cinerea*, and enhanced elf18-triggered immunity against *Pst*. Consistent with a negative role of ERF19 in PTI, MAMP-mediated growth inhibition was respectively weakened or augmented in lines overexpressing *ERF19* or expressing *ERF19-SRDX*. Moreover, we demonstrate that the transcriptional repressor Novel INteractor of JAZ (NINJA) associates with and represses the function of ERF19. Our work reveals ERF19 as a key player in a buffering mechanism to avoid defects imposed by over-activation of PTI and a potential role for NINJA in fine-tuning ERF19-mediated regulation.

## INTRODUCTION

Plants have adopted sophisticated defense mechanisms to fight off invading pathogens. Initiation of plant defense responses relies on the recognition of non-self organisms. Plants utilize pattern recognition receptors (PRRs) as the first line of surveillance to detect incoming threat posed by pathogens. Plant PRRs perceive microbe-associated molecular patterns (MAMPs), which are molecular structures conserved among microbes and crucial for the survival of microbes (Macho and Zipfel, 2014; Zipfel, 2014). For example, bacterial MAMP flg22, the active epitope of flagellin, is recognized by the PRR FLAGELLIN SENSING2 (FLS2)(Felix et al., 1999; Gómez-Gómez and Boller, 2000), and EF-Tu RECEPTOR (EFR) recognizes the conserved peptide elf18 derived from bacterial EF-Tu, which is one of the most abundant proteins in bacteria (Kunze et al., 2004; Zipfel et al., 2006). The fungal MAMP chitin, an important constituent of fungal cell walls (Silipo et al., 2010), is perceived by CHITIN ELICITOR RECEPTOR KINASE1 (CERK1) and LYSM-CONTAINING RECEPTOR-LIKE KINASE 5 (LYK5) (Miya et al., 2007; Wan et al., 2008; Cao et al., 2014). MAMP recognition induces pattern-triggered immunity (PTI), restricting the incursion and proliferation of potential pathogens (Boller and Felix, 2009; Schwessinger and Ronald, 2012; Newman et al., 2013).

Activation of PTI involves massive transcriptional reprogramming to mount defense responses against invading pathogens (Bigeard et al., 2015; Tsuda and Somssich, 2015; Garner et al., 2016; Birkenbihl et al., 2017a). General PTI responses include reinforcement of cell wall through deposition of callose and production of defense-related proteins (Boller and Felix, 2009). Pathogenesis-related (PR) proteins and plant defensins (PDF) represent two major classes of defense-related proteins with diverse antimicrobial activities (Thomma et al., 2002; van Loon et al., 2006). In *Arabidopsis*, *PR1* and *PR2* are induced after inoculation of the hemi-biotrophic bacterium *Pseudomonas syringae* pv. *tomato* DC3000 (*Pst*) and are marker genes for flg22 and elf18 treatments (Lu et al., 2009; Xiao and Chye, 2011; Nomura et al., 2012;), whereas *PDF1.2* and *PDF1.3*, which are induced by the necrotrophic fungus *Botrytis cinerea*, serve as potential markers for chitin elicitation (Pieterse et al., 2009; Pieterse et al., 2012; Meng et al., 2013).

Activation of plant immunity requires high expense of energy, and excessive immune responses reduce plant fitness, hampering plant growth and survival (Bolton, 2009; Katagiri and Tsuda, 2010; Kim et al., 2014). Transcription factors (TFs) lie at the heart of transcriptional reprograming, and the ethylene response factor (ERF) TF family plays a key role in orchestrating the balance of defense outputs (Huang et al., 2016; Jin et al., 2017). Perturbation of key immune regulators may tip the balance and lead to catastrophic growth retardation. For example, direct activation of ERF6 enhances *Arabidopsis* resistance to *B. cinerea* and induces constitutive activation of defense genes (Meng et al., 2013). However, these plants exhibit a severe dwarf phenotype, which might be the result of strong defense activation (Meng et al., 2013).

In order to maintain appropriate levels of defense activation, TFs that negatively regulate immunity need to work in concert with defense-activating TFs. For example, the pathogen-induced *ERF4* (*ERF078*) and *ERF9* (*ERF080*) negatively regulate *Arabidopsis* resistance against fungal pathogens and activation of *PDF1.2* (McGrath et al., 2005; Maruyama et al., 2013). In addition, transcriptional activities of TFs are modulated in a post-translational manner to ensure timely activation or repression of immune signaling (Licausi et al., 2013). Typically, ETHYLENE INSENSITIVE 3 (EIN3) transactivates *ERF1* (*ERF092*), but the transactivation function of EIN3 is repressed in the presence of JASMONATE ZIM-DOMAIN 1 (JAZ1) (Zhu et al., 2011). Notably, JAZ1 interacts with EIN3 and recruits the transcriptional repressor Novel Interactor of JAZ (NINJA) with TOPLESS (TPL) or TPL-related proteins (TPRs) (Pauwels et al., 2010; Zhu et al., 2011). EIN3-mediated activation of *ERF1* is de-repressed when JAZ1 is degraded upon accumulation of jasmonic acid (JA) that occurs after pathogen attack (De Vos et al., 2005; Chini et al., 2007; Zhu et al., 2011). JAZ1-imposed repression on EIN3 ensures that *ERF1* and ERF1-targeted defense genes such as *PDF1.2* are not induced in the absence of pathogen invasion (Pieterse et al., 2012).

While there are increasing reports showing that ERFs are involved in plant defense, whether ERFs directly participate in the regulation of PTI remains unclear. Here we report that the pathogen-and MAMP-induced *ERF19* plays a negative role in *Arabidopsis* immunity against both fungal and bacterial pathogens. Notably, overexpression of *ERF19* or repression of ERF19 function through expression of the chimeric dominant repressor *ERF19-SRDX* leads to respectively decreased and increased PTI responses. Our data further suggest that ERF19 functions as a modulator in MAMP-mediated growth inhibition and serves as a buffering mechanism to prevent detrimental effects of excessive PTI. Moreover, our biochemical and genetic approaches showed that NINJA associates with and represses the function of ERF19, suggesting another layer of control over PTI activation. Collectively, our functional studies on ERF19 provide evidences about an ERF involved in the regulation of PTI and new insights into the dynamic regulation of PTI.

## RESULTS

### Overexpression of *ERF19* Enhances *Arabidopsis* Susceptibility to Pathogens

To identify TFs involved in the regulation of *Arabidopsis* defenses against the fungal pathogen *B. cinerea*, we designed a screen to evaluate the resistance of *Arabidopsis* from the *At*TORF-Ex collection (Weiste et al., 2007; Wehner et al., 2011) to *B. cinerea.* Notably, we found a transgenic line overexpressing *ERF19/ERF019* (At1g22810, HA-ERF19) that developed increased disease lesions after drop-inoculation with *B. cinerea* spores (Supplemental Figures 1A-C). To ensure that the observed increased susceptibility phenotype of the HA-ERF19 line to *B. cinerea* was not due to multiple transformation events (Weiste et al., 2007), we generated additional *Arabidopsis* lines expressing the coding sequence (CDS) of *ERF19* fused with *GFP* under the control of the Cauliflower Mosaic Virus 35S (CaMV 35S) promoter in the Col-0 background. Two independent lines (ERF19-OE1 and -OE2), expressing high levels of *ERF19* mRNA and ERF19-GFP proteins (Supplemental Figures 2A and B), were selected for further analyses. Confirming the increased susceptibility to *B. cinerea* observed in HA-ERF19 (Supplemental Figures 1B and C), ERF19-OE1 and -OE2 developed larger disease lesions than Col-0 after *B. cinerea* drop-inoculation (Figure 1A). In addition to ERF19-OEs, we generated transgenic lines expressing the CDS of *ERF19-GFP* fusion under the control of the beta-Estradiol (β-Est)-inducible XVE system (ERF19-iOEs). Overexpression of *ERF19* and ERF19-GFP was β-Est dependent (Supplemental Figures 2C and D). Notably, increased susceptibility to *B. cinerea* was observed in ERF19-iOEs treated with β-Est but not in mock controls treated with DMSO (Supplemental Figure 3). Importantly, β-Est treatment did not alter Col-0 resistance against *B. cinerea* as compared to the DMSO-treated control (Supplemental Figure 3), indicating that the increased susceptibility to *B. cinerea* in ERF19-iOEs is specifically linked to overexpression of *ERF19* rather than to the β-Est treatment. In summary, our phenotypic analyses on HA-ERF19, ERF19-OEs, and ERF19-iOEs show that overexpression of *ERF19* enhances *Arabidopsis* susceptibility to *B. cinerea*. Confirming earlier work (Scarpeci et al., 2016), the rosette leaves of 5-week-old ERF19-OEs exhibited different degrees of inward curling, and the rosette biomass of ERF19-OEs was smaller than wild-type Col-0 (Supplemental Figures 4A and B). However, unlike ERF19-OEs, the rosettes of ERF19-iOE and Col-0 plants were indistinguishable when grown in laboratory conditions (Supplemental Figure 4C). This observation suggests that the enhanced susceptibility phenotype to *B. cinerea* observed in ERF19-OEs (Figure 1A), is not linked with the altered growth phenotype of these OE lines.

**Figure 1.**
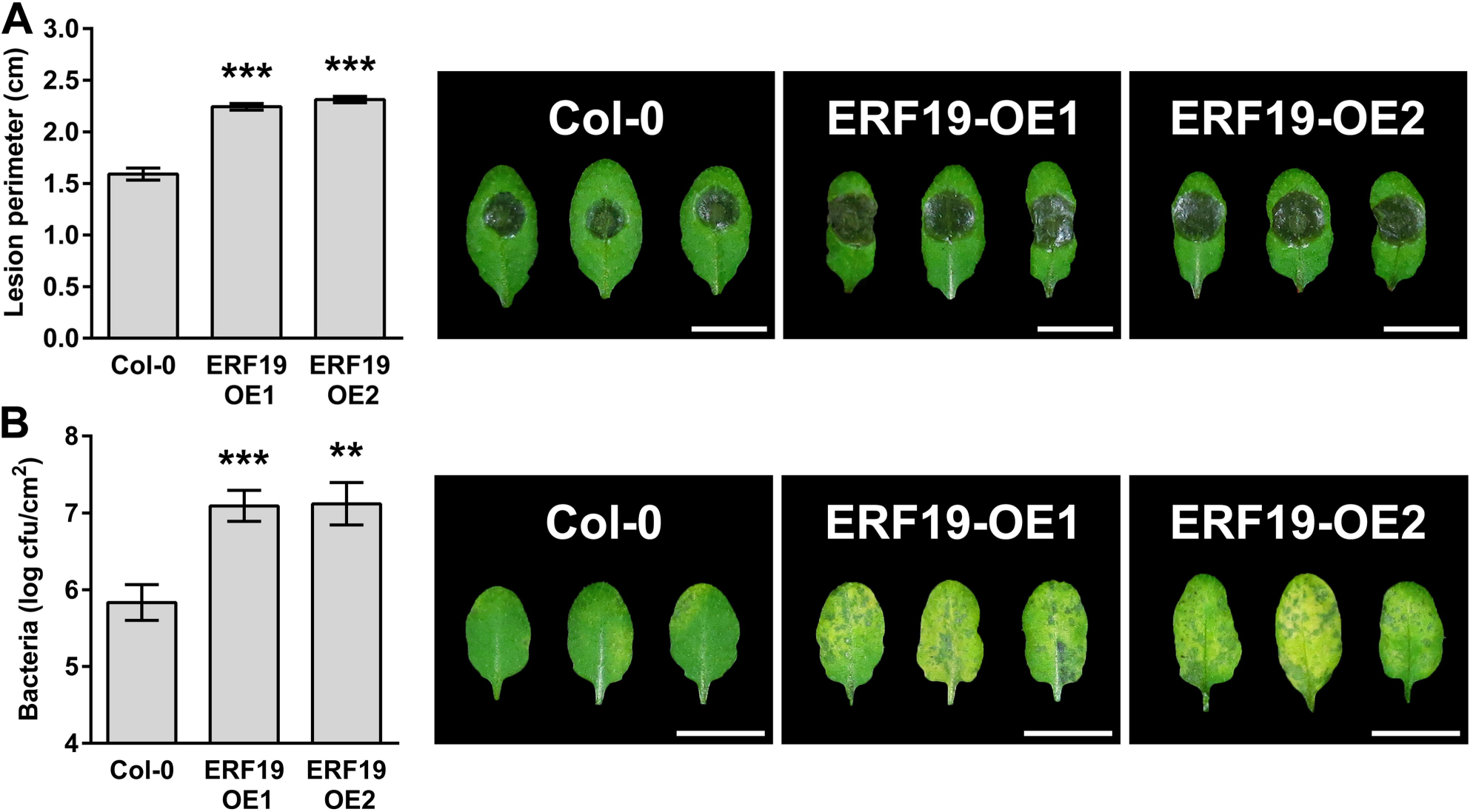
ERF19-OEs are hypersusceptibile to *B. cinerea* and *Pst*. (A) *B. cinerea*-mediated lesions. Leaves of 5-week-old ERF19-OEs droplet-inoculated with 8 μL of *B. cinerea* spore suspension (10^5^ spores mL^-1^ in 1/4 PDB). Disease symptoms were photographed and lesion perimeters were measured at 3 days post inoculation (dpi). Data represent average ± standard error (SE) of at least 72 lesion perimeters pooled from 3 independent experiments each with at least 6 plants per line. Asterisks indicate a significant difference to Col-0 based on a *t* test (\*\*\**P* < 0.001). (B) *Pst* growth and symptoms. Five-week-old plants were dip-inoculated with 10^6^ cfu mL^-1^ *Pst*, and symptoms were photographed at 3 dpi. Bacterial populations in the leaves were evaluated at 2 dpi. Values represent average ± SE from 3 independent experiments pooled each with 5 plants per line (n = 15). Asterisks indicate a significant difference to Col-0 based on a *t* test (\*\**P* < 0.01; *** *P* < 0.001).

**Figure 2.**
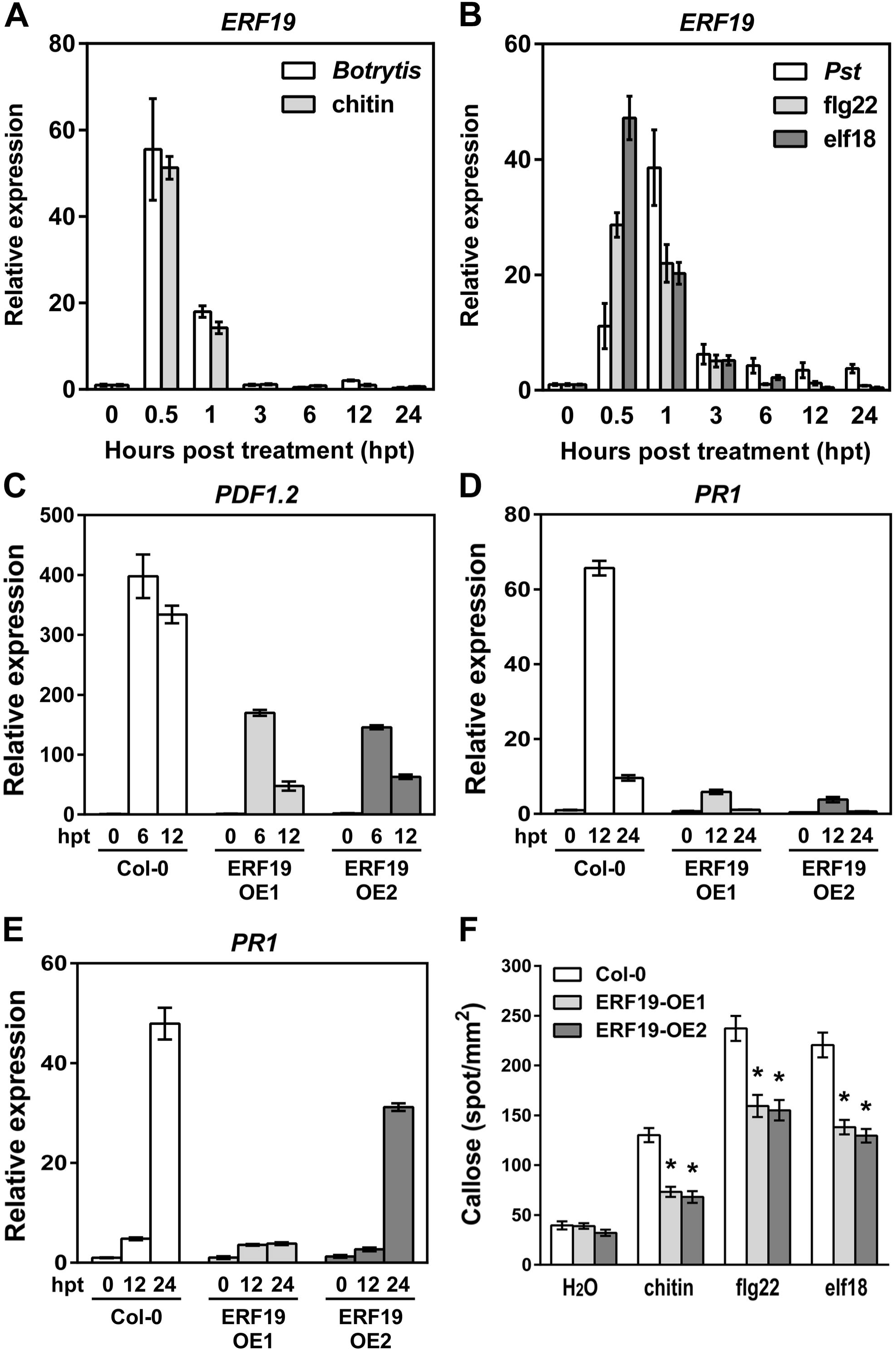
ERF19 is involved in PTI. (A) Time course expression of *ERF19* after inoculation with *B. cinerea* or treatment with chitin. Twelve-day-old seedlings were inoculated with a suspension of 5 × 10^5^ *B. cinerea* spores mL^-1^ or treated with 200 μg mL^-1^ chitin. Samples were collected at indicated time points, and *ERF19* expression was determined by qRT-PCR. After normalization with *UBQ10*, *ERF19* expression levels were compared to time 0 (defined value of 1). Error bars are SD from 3 technical replicates. Representative data are shown from one independent experiment repeated at least 3 times with similar results. (B) Time course expression of *ERF19* after inoculation with *Pst* or after treatment with flg22 or elf18. Twelve-day-old seedlings were inoculated with 10^7^ cfu mL^-1^ *Pst*, or treated with 100 nM flg22, or 100 nM elf18, and samples were collected at indicated time points. Analysis of *ERF19* expression was performed and presented as in (A). (**C** to **E**) Activation of PTI marker genes in ERF19-OEs. Chitin-induced *PDF1.2* (C), flg22-induced *PR1* (D), and elf18-induced *PR1* (E) in ERF19-OEs were determined by qRT-PCR. Twelve-day-old seedlings were treated with 200 μg mL^-1^ chitin, 1 μM flg22, or 1 μM elf18. Samples were collected at indicated time points, and analyses of PTI marker genes were performed and presented as in (A). (**F**) MAMP-induced callose deposition in ERF19-OEs. Fourteen-day-old seedlings were treated with deionized water (mock control), 200 μg mL^-1^ chitin, 100 nM flg22, or 100 nM elf18, and samples were collected 24 h later for aniline blue staining. Data represent the average numbers of callose deposits per square mm ± SE pooled from 4 independent experiments each with at least 6 biological repeats (n > 24). Asterisks denote values significantly different from respective Col-0 controls based on a *t* test. (\**P* < 0.01).

**Figure 3.**
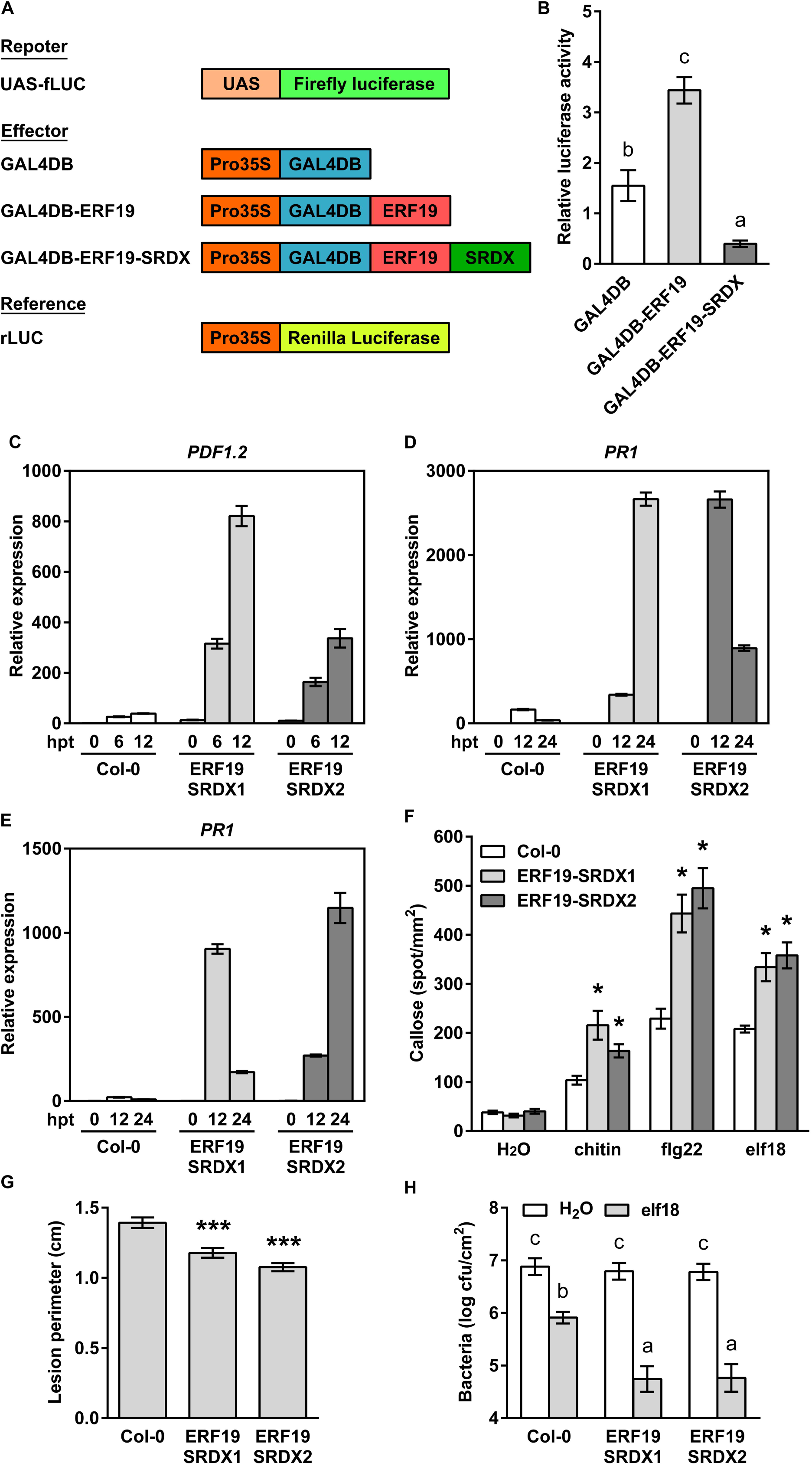
Expression of the dominant repressor *ERF19-SRDX* enhances PTI. (**A**) Schematic diagrams of reporter, effector, and reference plasmids used in the PTA assay (**B**) PTA assay. Relative luciferase activities were evaluated in *Arabidopsis* protoplasts co-transfected with the reporter plasmid (UAS:fLUC), the effector plasmids (35S:GAL4DB, 35S:GAL4DB-ERF19, or 35S:GAL4DB-ERF19-SRDX), and a calibrator plasmid encoding *rLUC*. Protoplasts transfected without the effector plasmids were used as a control (no effector). All the values were normalized to the rLUC activity and were relative to the values of the no effector control. Values are means ± SE of 4 independent experiments (n = 4). Different letters denote significant differences (*P* < 0.01) between groups based on a one-way ANOVA. (**C** to **E**) Activation of PTI marker genes in ERF19-SRDXs. Chitin-induced *PDF1.2* (C), flg22-induced *PR1* (D), and elf18-induced *PR1* (E) in ERF19-SRDXs were determined by qRT-PCR. Twelve-day-old seedlings were treated with 200 μg mL^-1^ chitin, 1 μM flg22, or 1 μM elf18. Samples were collected at indicated time points, and *UBQ10* was used for normalization. Relative gene expression levels were compared to Col-0 at time 0 (defined value of 1). Error bars indicate the SD for 3 technical replicates. Representative data are shown from one independent experiment repeated at least 3 times with similar results. (**F**) MAMP-induced callose deposition in ERF19-SRDXs. Fourteen-day-old seedlings were treated with deionized water (mock control), 200 μg mL^-1^ chitin, 100 nM flg22, or 100 nM elf18, and samples were collected 24 h later for aniline blue staining. Data represent the average numbers of callose deposits per square mm ± SE pooled from 3 independent experiments each with at least 6 biological repeats (n > 24). Asterisks denote values significantly different from respective Col-0 controls based on a *t* test. (\**P* < 0.01). (**G**) *B. cinerea*-mediated lesions in ERF19-SRDXs. Leaves of 5-week-old plants were droplet-inoculated with 8 μL of *B. cinerea* spores (10^5^ spores mL^-1^ in 1/4 PDB). Lesion perimeters were measured at 3 days post inoculation (dpi). Data represent average ± SE of 138 lesion perimeters (n = 138) pooled from 4 independent experiments each with at least 6 plants per line. Asterisks indicate a significant difference to Col-0 based on a *t* test (\*\*\**P* < 0.001). (**H**) *Pst* growth in ERF19-SRDXs. Five-week-old plants were syringe-infiltrated with H_2_ O or 10 nM elf18 6 h before syringe-infiltration with 10^6^ cfu mL^-1^ *Pst*. Bacterial populations in the leaves were evaluated at 2 dpi. Values represent average ± SE from 3 independent experiments pooled each with 3 plants per line (n = 9). Different letters denote significant differences between groups based on a two-way ANOVA (*P* < 0.01).

**Figure 4.**
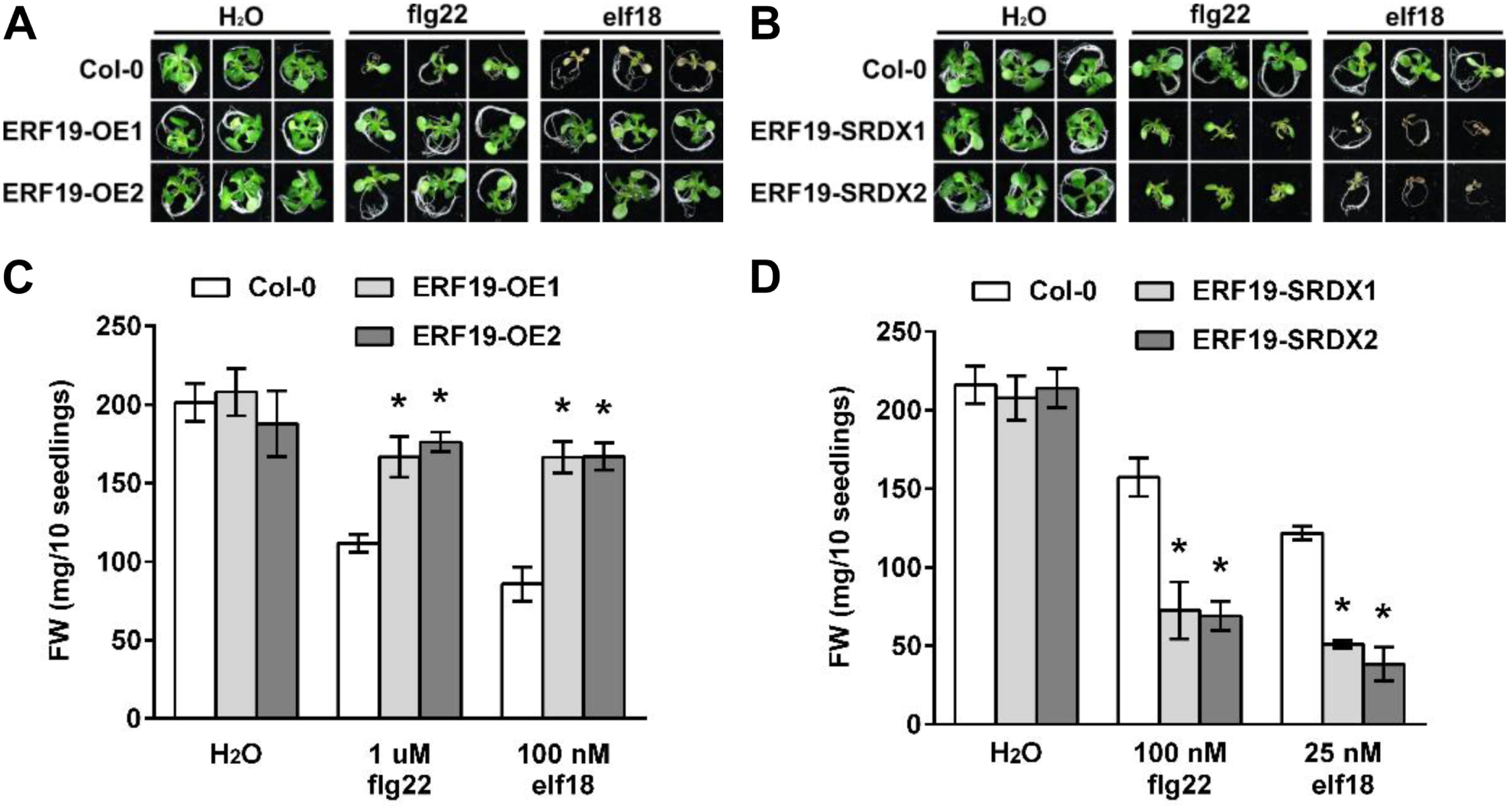
ERF19 negatively regulates MAMP-mediated growth inhibition. (**A** and **B**) Growth phenotypes. Five-day-old seedlings were grown in liquid 1/2 MS supplemented with 1 μM flg22 or 100 nM elf18 (A), or 100 nM flg22 or 25 nM elf18 (B). Seedlings were photographed 10 days after incubation. Scale bars represent 1 cm. (**C** and **D**) Fresh weight. Data represent the average fresh weight of 10 seedlings ± SE from 3 independent experiments (n = 3). Asterisks indicate a significant difference to respective Col-0 controls based on a *t* test (\**P* < 0.05).

To further dissect the role of ERF19 in *Arabidopsis* resistance to microbial pathogens, ERF19-OEs and Col-0 plants were dip-inoculated with *Pst*, and bacterial growth and disease symptoms were assessed 2 days later. ERF19-OEs harbored 10 times more bacteria and exhibited increased disease symptoms (Figure 1B), indicating that ERF19-OEs were hypersusceptible to *Pst* bacteria. Collectively, these data suggest that overexpression of *ERF19* in *Arabidopsis* induces hypersusceptibility to both fungal and bacterial pathogens.

### PTI Responses are Down-Regulated in *ERF19* Overexpression Lines

To further evaluate the role of ERF19 in *Arabidopsis* innate immunity, we monitored the expression of *ERF19* in Col-0 seedlings after inoculation with *B. cinerea* spores or treatment with the fungal MAMP chitin over a 24-hour period. *ERF19* transcripts were up-regulated by *B. cinerea* spores or chitin within half an hour, and steadily declined at later time points (Figure 2A). These results are consistent with previous reports showing that *ERF19* is rapidly induced by chitin and chitin derivatives (Ramonell et al., 2005; Libault et al., 2007; Fakih et al., 2016). Signaling pathways of phytohormones such as salicylic acid (SA), JA, and ethylene (ET) are important for transcriptional regulation of immune regulators (Pieterse et al., 2009; Pieterse et al., 2012). To further dissect the regulation of chitin-induced *ERF19*, we examined the expression of *ERF19* after chitin treatment in *npr1-1*, *coi1-16*, and *ein2-1* mutants, which are defective in SA, JA, and ET signaling pathways, respectively (Guzman and Ecker, 1990; Cao et al., 1994; Ellis and Turner, 2002). Chitin-induced *ERF19* transcripts in *ein2-1*, *npr1-1*, and *coi1-16* were similar to wild-type Col-0 within 1 hour post treatment (Supplemental Figures 5A and B). These data suggest that rapid induction of *ERF19* by chitin is unaffected when SA, JA, or ET signaling is impaired.

**Figure 5.**
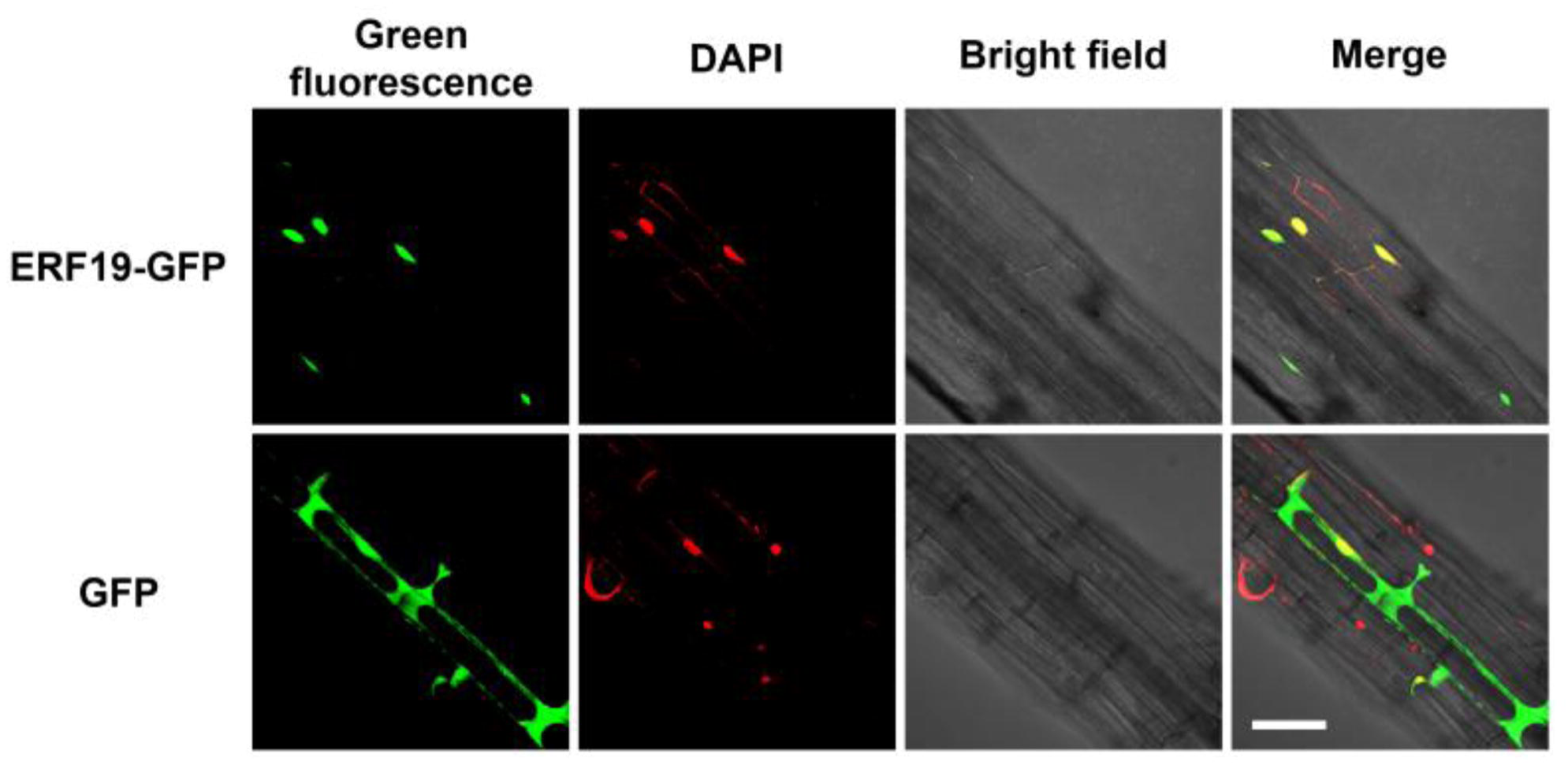
Subcellular localization of ERF19-GFP. Pictures were taken from seedlings of 12-day-old ERF19-iOE1 treated with 20 μM β-Est for 24 h and 35S:GFP transgenic lines. DAPI staining was used to determine the position of nuclei. Strong green fluorescence of ERF19-GFP was co-localized with the DAPI-stained nuclei. Scale bar represents 10 μm.

Since overexpression of *ERF19* induced hypersusceptibility to *Pst* bacteria, we also monitored the expression of *ERF19* in Col-0 seedlings after inoculation with *Pst*, or after treatment with the bacterial MAMPs flg22, or elf18. Similarly to *B. cinerea* spores or chitin, inoculation with *Pst*, or treatments with flg22 or elf18 transiently up-regulated *ERF19* for 1 hour, but *ERF19* transcripts steadily declined afterwards (Figure 2B).

Plants utilize PTI as a defense mechanism to ward off diverse pathogens (Boller and Felix, 2009; Huang and Zimmerli, 2014), and perturbation of PTI compromises plant defense against both fungal and bacterial pathogens. (Tsuda et al., 2009; Kim et al., 2014). Since ERF19-OEs showed an increased susceptibility to both *B. cinerea* and *Pst* and that *ERF19* is up-regulated by fungal and bacterial MAMPs, we evaluated whether ERF19 is involved in PTI. Towards this goal, we monitored the expression of PTI maker genes in ERF19-OEs and Col-0 after MAMP treatments. Notably, transcripts of chitin-induced *PDF1.2* and *PDF1.3* were lower in ERF19-OEs than in Col-0 (Figure 2C and Supplemental Figure 6A). Similarly, *PR1* and *PR2* expression induced by flg22 or elf18 were lower in ERF19-OEs than in Col-0 (Figures 2D and E; Supplemental Figures 6B and C), indicating that the up-regulation of these PTI marker genes is repressed when *ERF19* is overexpressed. We next measured callose deposition, a PTI output activated by fungal and bacterial MAMPs (Millet et al., 2010; Shinya et al., 2014), in ERF19-OEs and Col-0. While the water-treated callose deposits were similar between ERF19-OEs and Col-0, callose deposition induced by chitin, flg22, or elf18 was significantly impaired in ERF19-OEs (Figure 2F). Taken together, these results demonstrate that overexpression of *ERF19* represses common PTI responses, and the reduced PTI may account for the increased pathogen susceptibility observed in ERF19-OEs.

**Figure 6.**
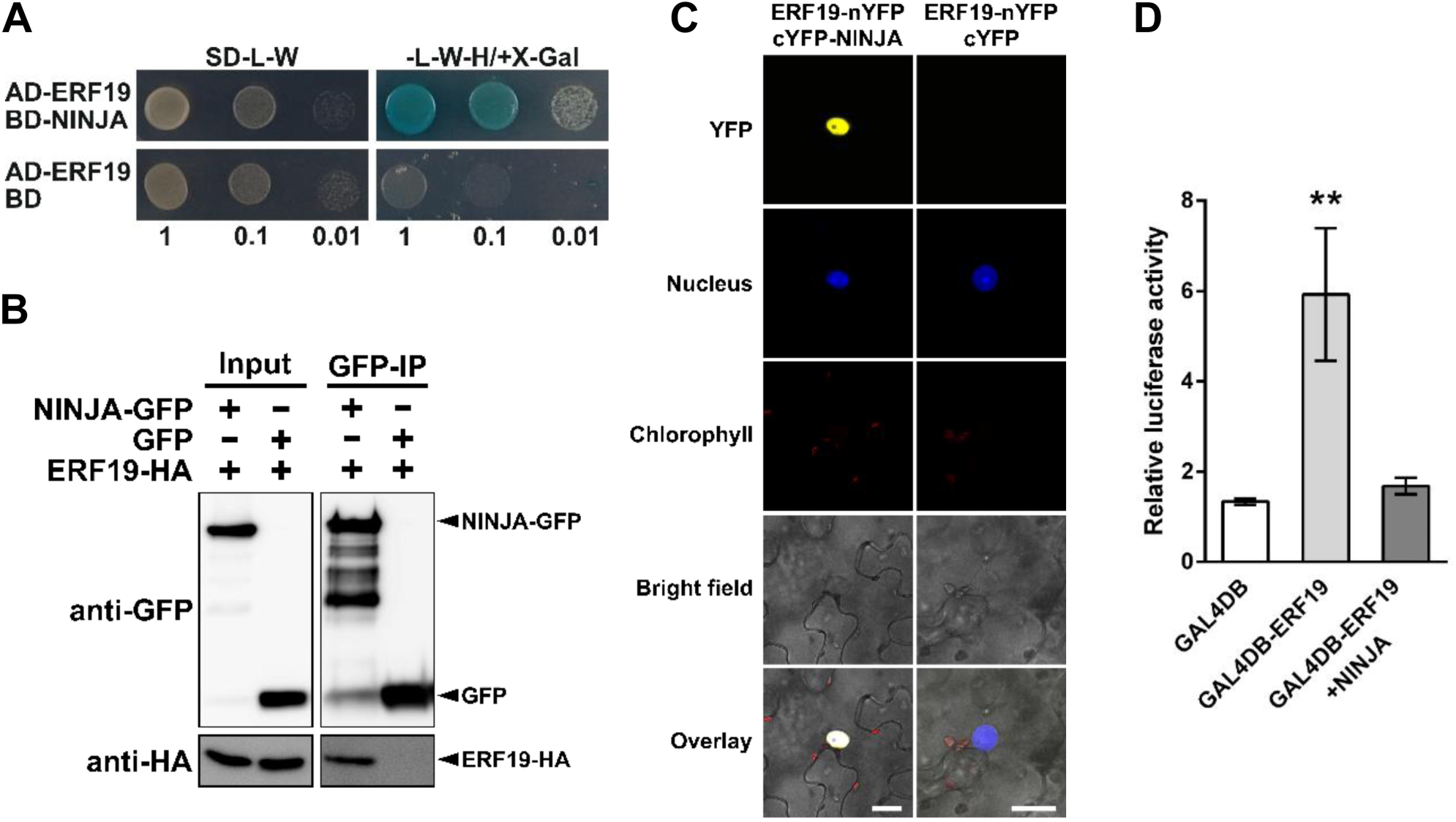
NINJA associates and represses the transcriptional activity of ERF19. (**A**) ERF19-NINJA association in Y2H assays. Ten-fold serial dilutions of yeasts expressing the indicated protein fusion to the activation domain (AD) or binding domain (BD) of GAL4 were plated on control (-L-W) or selective (-L-W-H/+X-α-Gal) SD media. Growth and blue staining of the colonies on selective SD medium indicate association between the two fusion proteins. (**B**) Analysis of ERF19-NINJA association by Co-IP. Total proteins from protoplasts expressing GFP or NINJA-GFP with ERF19-HA_3_ were immunoprecipitated (IP) with anti-GFP antibodies. Total proteins before (input) and after IP (GFP-IP) were immunoblotted with anti-GFP and anti-HA antibodies. Similar results were obtained from 3 independent experiments. (**C**) BiFC analysis of ERF19-NINJA association. *N. benthamiana* plants were co-transformed with the indicated split YFP constructs and a nuclear marker construct carrying NLS-mCherry-mCherry-NLS. YFP fluorescence (yellow), nucleus (blue), chlorophyll autofluorescence (red), bright field, and overlay images are shown. This experiment was performed at least 3 times with similar results. Scale bars represent 20 μm. (**D**) PTA assay. Relative luciferase activities of *Arabidopsis* protoplasts co-transfected with the reporter plasmid (UAS:fLUC), the effector plasmids (35S:GAL4DB, 35S:GAL4DB-ERF19, or 35S:GAL4DB-ERF19 and 35S:NINJA), and a normalization plasmid encoding *rLUC*. All the values were normalized to the rLUC activity and were relative to the values of the no effector control. Data are means ± SE of 7 independent experiments (n = 7). Asterisks indicate significant differences compared to the 35S:GAL4DB control (\*\**P* < 0.01).

### Expression of a Dominant Negative *ERF19-SRDX* Transgene Enhanced *Arabidopsis* PTI Responses

To perform a loss-of-function analysis of *ERF19*, we generated transgenic lines expressing a chimeric ERF19, as no mutants with insertions at the CDS or the proximal promoter region of *ERF19* are available. Transgenic expression of chimeric TFs, which exhibit dominant-negative function of the TFs, is a commonly used strategy for loss-of-function analysis of TFs (Mitsuda and Ohme-Takagi, 2009). Moreover, this approach overcomes the problem of functional redundancy, which might occur in single gene knockout or knockdown methods (Mitsuda and Ohme-Takagi, 2009). We first examined the transcriptional activity of ERF19 by using protoplast transactivation (PTA) assays based on the GAL4/upstream activation sequence (UAS) and dual-luciferase reporter system. In *Arabidopsis* protoplasts, expression of ERF19 fused to GAL4 DNA binding domain (GAL4DB) showed higher luciferase activity than expression of GAL4DB alone (Figures 3A and B), suggesting that ERF19 acts as a transcription activator. For loss-of-function study of activators, plant-specific EAR-motif repression domain (SRDX) are fused to activators to create chimeric repressors (Hiratsu et al., 2003; Mitsuda et al., 2011). Importantly, PTA assays revealed that the fusion of SRDX to ERF19 successfully converted the activator feature of ERF19 into a repressor (Figure 3B), indicating that the chimeric repressor ERF19-SRDX would suffice for loss-of-function analysis.

To further assess the biological function of ERF19-SRDX, we expressed the *ERF19* genomic sequence, consisting of the intergenic promoter region (−1 to −1535 bp), 5’ untranslated region, and CDS of *ERF19* fused to *SRDX* CDS, in Col-0 to generate ERF19-SRDX lines. The use of native promoter of *ERF19* better reflects the biological function of ERF19-SRDX than a constitutive promoter (Mitsuda and Ohme-Takagi, 2009). Two independent lines of ERF19-SRDXs, of which the transgene *ERF19-SRDX* was chitin responsive, were selected for further analyses (Supplemental Figure 7A). Unlike ERF19-OEs, the rosettes of ERF19-SRDXs were undistinguishable from Col-0 wild-type (Supplemental Figure 7B). To confirm the role of ERF19 in PTI and pathogen resistance, we first analyze MAMP responses of ERF19-SRDXs. Notably, chitin-induced *PDF1.2* and *PDF1.3*, flg22-induced *PR1*, and elf18-induced *PR1* and *PR2*, were higher in ERF19-SRDXs than in Col-0 (Figures 3C to E; Supplemental Figures 7C and E). Surprisingly, despite enhanced expression of flg22-induced *PR1*, ERF19-SRDXs showed wild-type expression levels of flg22-induced *PR2* (Supplemental Figure 7D). We also tested MAMP-induced callose deposition, and in contrast to ERF19-OEs, MAMP-induced callose deposits were higher in ERF19-SRDXs than in Col-0 (Figure 3F). Together, these results suggest that transgenic expression of *ERF19-SRDX* enhances *Arabidopsis* PTI responses.

**Figure 7.**
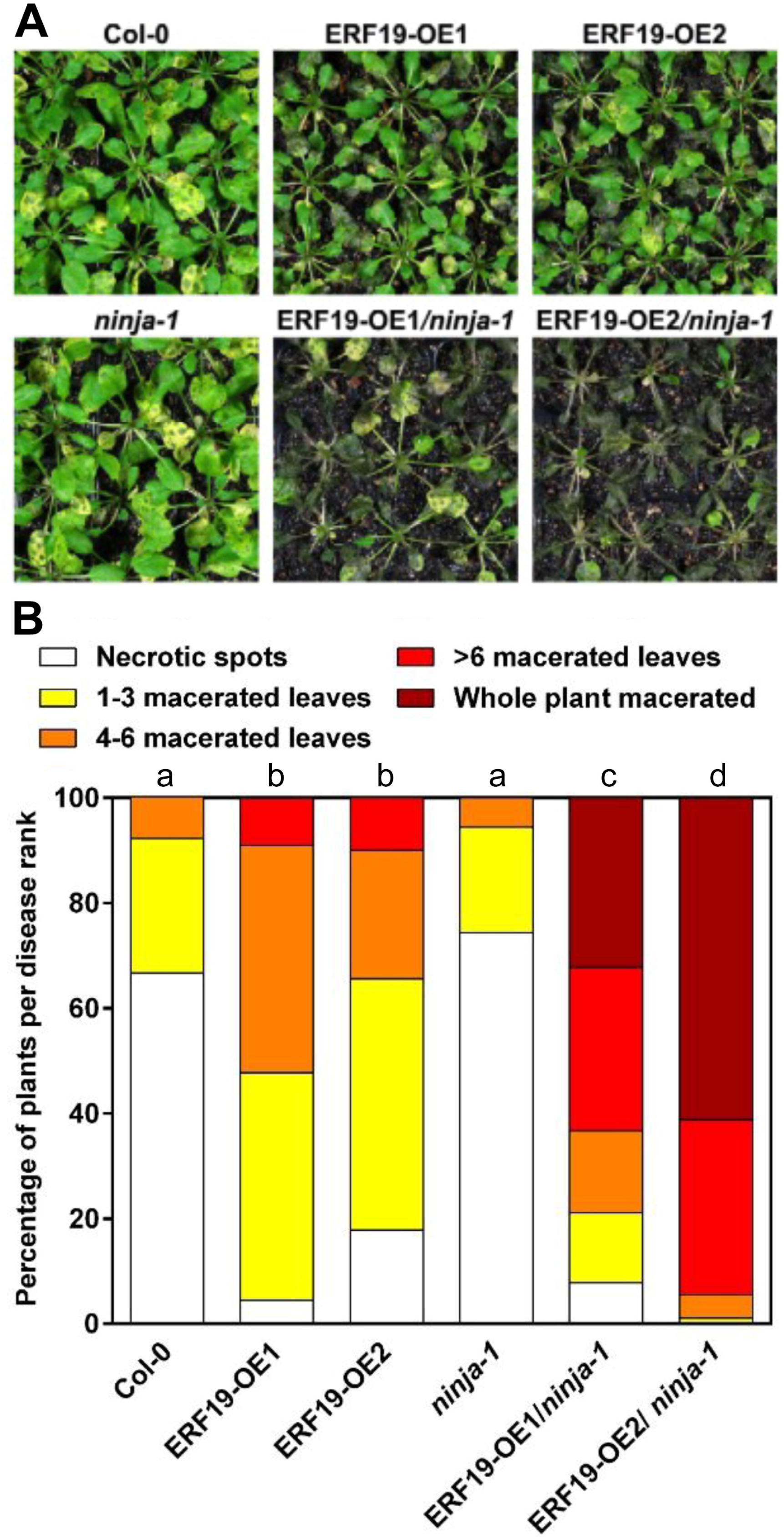
Hypersusceptibility to *B. cinerea* by *ERF19* overexpression is enhanced in *ninja-1*. (**A** and **B**) *B. cinerea* resistance in transgenic lines overexpressing *ERF19*. Four-week-old plants were spray-inoculated with a *B. cinerea* spore suspension (10^5^ spores mL^-1^ in 1/4 PDB). Symptoms were photographed (A) and disease ranks were determined (B) at 5 dpi. Data in (B) represent 90 biological replicates (n = 90) pooled from 3 independent experiments. The distribution of the disease rank proportions among the lines was analyzed using the chi-squared test. Groups that do not share a letter are significantly different in the distribution of disease ranks (*P* < 0.01).

In ERF19-OEs, the enhanced susceptibility to fungal and bacterial pathogens was correlated with reduced PTI responses. We thus hypothesized that the heightened PTI activation in ERF19-SRDXs will confer pathogen resistance. Indeed, ERF19-SRDXs exhibited smaller disease lesions than Col-0 after *B. cinerea* infection (Figure 3G), indicating ERF19-SRDXs were more resistant to *B. cinerea* than Col-0 wild-type plants.

However, Col-0 and ERF19-SRDXs developed similar disease symptoms after *Pst* infection (Supplemental Figure 7F), suggesting ERF19-SRDXs are as susceptible as Col-0 to *Pst* bacteria. *Pst* is an aggressive pathogen armed with a plethora of virulence factors that suppress host immunity (Dou and Zhou, 2012; Xin and He, 2013). We thus speculated that the observed ERF19-SRDXs wild-type susceptibility to *Pst* is associated with subversion of the activation of host PTI. To highlight the role of ERF19-SRDX in PTI-mediated defense against *Pst*, we therefore activated plant PTI by treatment with 10 nM of the MAMP elf18 prior to *Pst* inoculation. In water-treated controls, bacterial growth was similar in Col-0 and ERF19-SRDXs (Figure 3H), confirming our previous observation that ERF19-SRDXs showed wild-type susceptibility to *Pst* (Supplemental Figure 7F). Strikingly, decrease of *Pst* growth by elf18 pretreatment was significantly stronger in ERF19-SRDXs than in Col-0 (Figure 3H), suggesting that elf18-induced resistance to *Pst* was enhanced in ERF19-SRDXs. Together, these results show that the expression of the dominant repressor *ERF19-SRDX* boost PTI responses and consequently, confers increased resistance to fungal and bacterial pathogens. In summary, our phenotypic analyses on ERF19-OEs and ERF19-SRDXs provide genetic evidences that ERF19 plays a negative role in the regulation of *Arabidopsis* PTI and defense towards pathogens.

### Perturbation of *ERF19* Disrupts MAMP-Mediated Growth Arrest

PTI costs plants high amount of energy, and thus the activation of PTI must be kept under tight control (Bolton, 2009; Katagiri and Tsuda, 2010; Kim et al., 2014). The fact that *ERF19* is induced upon elicitation by pathogens or MAMPs, and that ERF19 negatively regulates PTI, prompted us to test whether ERF19 protects plant from over-activation of PTI. For this purpose, we evaluated through *ERF19* gain-and loss-of-function analyses whether ERF19 alters plant sensitivities toward flg22-and elf18-mediated growth arrest, a well-documented feature of PTI (Gómez-Gómez and Boller, 2000; Zipfel et al., 2006; Ranf et al., 2011). Although 5-week-old Col-0 and ERF19-OEs had different growth habits (Supplemental Figure 4A), in our experimental conditions, water control seedlings of 15-day-old Col-0 and ERF19-OEs had similar fresh weight (Figures 4A and C). Remarkably, treatment with flg22 or elf18 profoundly inhibited the growth of Col-0 seedlings (Figures 4A and C). However, the MAMP-mediated growth inhibition effect was significantly lower in ERF19-OEs (Figures 4A and C). By contrast, treatment with flg22 or elf18 at low concentration severely inhibited the growth of ERF19-SRDXs whereas the growth of Col-0 was affected to a lesser degree (Figures 4B and D). Therefore, decreased and increased sensitivities to MAMP-mediated growth inhibition parallel the reduced and enhanced PTI in ERF19-OEs and ERF19-SRDXs (compare Figures 2 and 3 with Figure 4). Together, these results suggest that ERF19 plays a crucial role in tuning down PTI to protect *Arabidopsis* from detrimental growth inhibition effects imposed by excessive PTI.

### ERF19 is a Nuclear TF

To determine the subcellular localization of ERF19, we took advantage of the high expression levels of ERF19-GFP in β-Est-treated ERF19-iOE1. Confocal microscope images revealed that strong GFP signals co-localized with DAPI-stained nuclei in the seedling roots of β-Est-treated ERF19-iOE1 (Figure 5), indicating that ERF19-GFP is enriched in the nucleus. On the other hand, roots of transgenic seedlings constitutively expressing GFP alone showed dispersed nuclear and cytoplasmic fluorescence (Figure 5).

### NINJA Associates With and Represses ERF19

The activities of TFs can be influenced via protein-protein interactions, and identification of TF-interacting proteins is thus crucial to resolve the complex regulation of TF-signaling networks (Licausi et al., 2013). We thus investigated whether ERF19 is regulated via protein-protein interaction. Our pilot experiments based on yeast two-hybrid (Y2H) assays showed that ERF19 was capable of associating with the transcriptional repressor NINJA *in vitro* (Figure 6A) (Pauwels et al., 2010). Supporting this observation, immunoprecipitation with GFP antibody in *Arabidopsis* protoplasts showed that ERF19-HA_3_ proteins could be pulled down along with NINJA-GFP proteins (Figure 6B), suggesting that ERF19 associates with NINJA *in planta*. In contrast, no ERF19-HA_3_ proteins were detected when GFP was used for immunoprecipitation (Figure 6B). In addition, bimolecular fluorescence complementation (BiFC) data showed that the reconstitution of YFP in the nucleus was observed when ERF19 fused to the N-terminus of YFP (ERF19-nYFP) and NINJA fused to the C-terminus of YFP (cYFP-NINJA) were co-expressed in the leaves of *Nicotiana benthamiana* (Figure 6C). Co-expression of ERF19-nYFP and cYFP alone did not show any yellow fluorescence (Figure 6C). Together, these data indicate that ERF19 associates with NINJA *in planta*.

NINJA represses the transactivation activities of TFs through connection with co-repressors such as TPL or TPRs (Pauwels et al., 2010). To further understand the rationale of ERF19-NINJA association, we tested via PTA analysis whether NINJA alters the transcriptional activity of ERF19. Co-transfection of GAL4DB-ERF19 with NINJA significantly repressed ERF19-activated luciferase activity (Figures 6D), suggesting that NINJA plays a negative role on the transcriptional activity of ERF19. To clarify the biological role of NINJA in ERF19 function, we employed a genetic approach by overexpressing *ERF19* in the *NINJA* loss-of-function mutant *ninja-1* (Acosta et al., 2013). Two independent lines (ERF19-OEs/*ninja-1*) overexpressing *ERF19* mRNA and ERF19-GFP proteins, with comparable expression levels to ERF19-OEs in the Col-0 background (Supplemental Figures 8A and B), were selected for phenotypic analysis. Interestingly, the *ninja-1* mutant appeared to have a long petiole phenotype when grown in our laboratory conditions, and ERF19-OEs/*ninja-1* showed dramatically reduced rosette sizes (Supplemental Figure 8C). It was difficult to evaluate *B. cinerea*-mediated disease lesions by droplet-inoculation on ERF19-OEs/*ninja-1* due to their small leaf sizes. The disease resistance of ERF19-OEs/*ninja-1* against *B. cinerea* was thus assessed through spray-inoculation and progression of *B. cinerea* was ranked according to disease symptoms. Remarkably, after spray-inoculation with *B. cinerea* spores, ERF19-OEs/*ninja-1* developed dramatic disease symptoms. Notably, most of the plants were heavily or completely macerated (Figure 7A and B). By contrast, ERF19-OE plants exhibited only several macerated leaves, and symptoms were less severe than ERF19-OEs/*ninja-1* (Figure 7A and B). While the majority of the Col-0 and *ninja-1* plants developed symptoms with necrotic spots, they showed the least severe symptoms of the lines tested (Figure 7A and B). These results indicate that loss of *NINJA* function strongly enhanced susceptibility to *B. cinerea* in *ERF19* overexpression lines. This observation further implies that NINJA represses the function of ERF19 in *Arabidopsis* immunity. In summary, our data based on biochemical and genetic approaches strongly suggest that NINJA associates and negatively regulates the function of ERF19.

**Figure 8.**
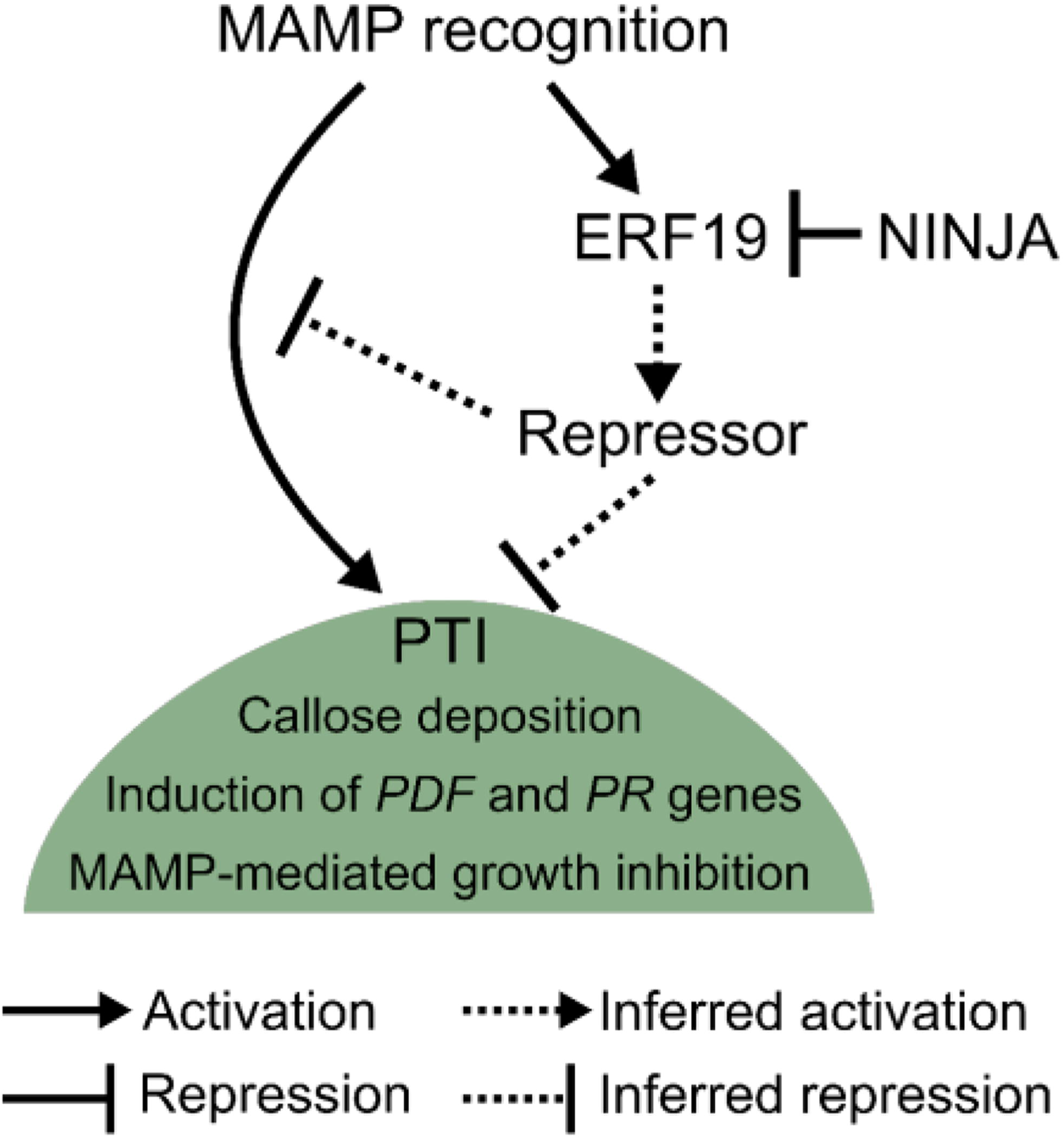
Proposed model for ERF19 and NINJA actions. Upon MAMP perception, *ERF19* is induced in parallel with PTI activation signals. Accumulation of ERF19 may transcriptionally induce repressors, which are likely to be involved in direct or indirect suppression of PTI signaling. PTI responses such as callose deposition, induction of *PDF* and *PR* genes, and MAMP-induced growth arrest are tuned-down by the ERF19-mediated pathway. The repressor NINJA provides another layer of control on PTI signaling through negative regulation of ERF19 function.

## DISCUSSION

### ERF19 Negatively Regulates PTI

*ERF19* was first identified as one of the genes highly induced by chitin (Libault et al., 2007) and was used as a marker for chitin-elicitation (Fakih et al., 2016). ERF19 is also involved in the regulation of plant growth, flowering time, senescence, and water-deficit stress (Scarpeci et al., 2016). Here we report that ERF19 functions as a negative regulator of PTI. Our gain-and loss-of-function analyses based on phenotypic studies of ERF19-OEs and ERF19-SRDXs revealed that ERF19 negatively regulates disease resistance against the fungus *B. cinerea* and *Pst* bacteria. Although ERF19-OEs exhibited curly leaves and reduced rosette size, the increased disease susceptibility of ERF19-OEs is likely not linked to the altered developmental habitus of *ERF19* overexpression. Indeed, we showed that ERF19-iOEs with appearance and morphology undistinguishable from wild-type Col-0 were also hypersusceptible to *B. cinerea* when *ERF19* overexpression was induced by β-Est. These observations suggest that plant growth is not the major determinant of ERF19-mediated susceptibility. In line with this argument, small size plants, as a result of overexpression of TFs, could either display increased or decreased resistance against pathogens (Chen and Chen, 2002; Xing et al., 2008; Tsutsui et al., 2009), further suggesting that plant growth habitus is not a decisive measure of plant resistance. Importantly, the altered *B. cinerea* and *Pst* resistance in ERF19-OEs and ERF19-SRDXs was correlated with an altered activation of PTI. PTI functions through common signaling pathways to transcriptionally activate defense responses against invading pathogens (Kim et al., 2014). The necrotrophic fungus *B. cinerea* and the hemi-biotrophic bacterium *Pst* are distinct microorganisms and therefore, the altered resistance to both pathogens observed may be a result of perturbations of a common sector in the PTI signaling network. Indeed, activation of MAMP-specific marker genes were repressed in ERF19-OEs and enhanced in ERF19-SRDXs, suggesting that ERF19 negatively regulates the PTI signaling network. In addition, *ERF19* was induced by fungal and bacterial MAMPs, and the diverse natures of these MAMPs further imply that ERF19 is a critical regulator in a common, general PTI signaling network. Furthermore, it is unlikely that ERF19 directly targets the PTI marker genes we tested, since the transient induction of *ERF19* was hours before the induction of these marker genes. Moreover, since ERF19 acted as a transcriptional activator when analyzed by PTA, PTI marker genes directly targeted by ERF19 should be up-regulated and not down-regulated as observed, by *ERF19* overexpression. In addition, the possibility that ERF19 acts as a repressor in a locus-specific manner (Lefstin and Yamamoto, 1998; Liu et al., 2015a) and directly represses the PTI marker genes is precluded, as the expression of the dominant repressor *ERF19-SRDX* resulted in enhanced up-regulation of the PTI marker genes tested. Based on these observations, we propose that the repression of PTI signaling by ERF19 is likely mediated through the activation of transcriptional repressors. These repressors in turn directly or indirectly repress PTI signaling pathways (Figure 8).

### Transcriptional Regulation of *ERF19*

Rapid and transient up-regulation of *ERF19* by pathogens and MAMPs may seem paradoxical, since ERF19 plays a negative role in PTI activation. In fact, positive and negative regulators of immunity work in concert to mount appropriate levels of defense responses (Couto and Zipfel, 2016). In line with this, *ERF4*, *ERF9*, rice *OsERF922*, and potato *StERF3* are induced by pathogens and function as negative regulators in plant immunity (McGrath et al., 2005; Liu et al., 2012; Maruyama et al., 2013; Tian et al., 2015). In addition, the *L-TYPE LECTIN RECEPTOR KINASE-V.5* (*LecRK-V.5*), which is induced specifically in stomatal guard cells by *Pst* and flg22, negatively regulates pathogen-and MAMP-induced stomatal closure, a common response of PTI (Melotto et al., 2006; Arnaud et al., 2012; Desclos-Theveniau et al., 2012). In addition, flg22-induced *WRKY18* and *WRKY40* act redundantly to negatively regulate flg22-triggered genes (Birkenbihl et al., 2017b). Collectively, these studies show that recognition of pathogens or MAMPs can transcriptionally induce negative regulators of immunity, which are necessary to buffer plant defense outputs.

The SA, JA, and ET pathways are suggested to play important roles to regulate up-regulation of pathogen-induced TFs. For example, expression of *ERF1* after *Fusarium oxysporum* f. sp. *conglutinans* inoculation depends on JA and ET signaling pathways and is independent of SA (Berrocal-Lobo and Molina, 2004). Similarly, *B. cinerea*-induced *ERF96* requires intact JA and ET pathways (Catinot et al., 2015). In contrast, JA and ET signalings negatively regulate *Pst*-induced *WRKY48* (Xing et al., 2008). By using appropriate mutants, we showed that rapid induction of *ERF19* by chitin was unaffected when SA, JA, and ET signalings were individually impaired. It is possible that SA, JA, and ET signalings act redundantly in the transcriptional control of chitin-(or MAMP-) induced *ERF19* so that loss-of-function of one pathway would be compensated by other functional signaling pathways. Another possibility is that the induction of *ERF19* by chitin (or MAMPs) is regulated in addition to or independently of SA, JA, and ET. Other cellular responses downstream of MAMP perception such as the activation of mitogen-activated protein kinase (MPK) cascades, ROS burst, and changes in cellular calcium levels also mediate transcriptional regulation of PTI (Torres, 2010; Reddy et al., 2011; Li et al., 2016). For example, while *ERF6* is moderately induced by JA and ET treatments (Moffat et al., 2012; Son et al., 2012), direct activation of MPK3/MPK6 is sufficient to substantially induce *ERF6* expression (Meng et al., 2013). A transcriptome analysis on catalase-deficient plants revealed that *ERF19* is up-regulated in response to high light-induced H_2_ O_2_ treatment (Vanderauwera et al., 2005). ROS burst is a conserved response after MAMP-elicitation (Boller and Felix, 2009; Torres, 2010), and the rapid and transient pattern of MAMP-induced ROS burst (Monaghan et al., 2014) is similar to the expression pattern of MAMP-induced *ERF19*. It is thus tempting to postulate that MAMP-induced ROS takes part in the transcriptional activation of *ERF19*.

### ERF19 Buffers MAMP-Induced Growth Inhibition

Plant growth and immunity are maintained at a fine balance to ensure plant survival. In the presence of invading pathogens, positive and negative regulators of immunity together tailor this balance to ensure appropriate levels of defense outputs. Exaggerated defense responses that tip the balance towards immunity can hamper plant growth and survival. For example, constitutive activation of ERF6 or overexpression of *ERF11* results in direct activation of defense genes, but these transgenic plants suffers from severe growth detriment (Tsutsui et al., 2009; Meng et al., 2013). In addition, overexpression of BRASSINOSTEROID INSENSITIVE1-ASSOCIATED RECEPTOR KINASE1 (BAK1), which interacts with FLS2 and EFR and acts as a co-receptor for flg22 and elf18 (Chinchilla et al., 2007; Roux et al., 2011), directly activates PTI in the absence of MAMPs. However, BAK1 overdose results in stunted growth, leaf necrosis, and decreased seed production (Dominguez-Ferreras et al., 2015). Similarly, the L-TYPE LECTIN RECEPTOR KINASE-VI.2 (LecRK-VI.2) associates with FLS2 and functions as a positive regulator of PTI (Singh et al., 2012; Huang et al., 2014). Plants with high expression of *LecRK-VI.2* show constitutive PTI responses but display a dwarf phenotype (Singh et al., 2012). On the other hand, loss of *BAK1-INTERACTING RECEPTOR-LIKE KINASE 1* (*BIR1*), a negative regulator of plant immunity, leads to constitutive activation of defense responses and cell death, which dramatically hampers plant growth (Gao et al., 2009). These studies illustrate that genetic disruption of crucial immune regulators can deleteriously affect plant growth. Although ERF19 functions as a negative regulator of PTI, unlike *bir1* mutant (Gao et al., 2009), *ERF19* loss-of-function lines ERF19-SRDXs showed wild-type growth under normal conditions and did not exhibit constitutive activation of PTI responses. The dominant repressor *ERF19-SRDX* was regulated by the native promoter of *ERF19*. This basal expression of *ERF19-SRDX* might thus be insufficient to trigger PTI activation. In spite of normal growth and basal PTI responses in ERF19-SRDXs, the effects of flg22-or elf18-induced growth inhibition was much more severe on ERF19-SRDXs than on Col-0, even at low concentration of flg22 or elf18. The high sensitivity of ERF19-SRDXs to MAMP-mediated growth arrest implies that in response to MAMPs, ERF19 acts as a buffering regulator of PTI to prevent exaggerated defense responses, which could negatively impact plant growth. In agreement with this, ERF19-OEs showed diminished growth inhibition imposed by high concentration of MAMPs. Taken together, our data pinpoint ERF19 as being part of an *Arabidopsis* PTI buffering mechanism to maintain the balance between growth and immunity upon MAMP recognition.

### NINJA Negatively Regulates ERF19

Post-translational regulation such as protein-protein interaction may alter the transcriptional activities of TFs (Licausi et al., 2013). For example, JAZ1 interacts with and represses the transcriptional function of EIN3 (Zhu et al., 2011). In addition, MYC2, a crucial TF regulating JA signaling, interacts with EIN3 and represses the transcriptional activity and DNA binding ability of EIN3. Conversely, EIN3 interacts with MYC2 and represses the transactivation function of MYC2 (Song et al., 2014; Zhang et al., 2014). JAZ proteins also play negative roles in regulating the function of MYC TFs through direct interaction (Chini et al., 2007; Pauwels and Goossens, 2011; Zhang et al., 2015). Such negative regulations are thought to modulate fine-tuning mechanisms to achieve rigorous transcriptional controls. NINJA was originally identified as the adaptor between JAZ proteins and the transcriptional co-repressors TPL and TPRs and was demonstrated to act as a negative regulator of JA signaling (Pauwels et al., 2010). Later studies showed that NINJA is also involved in the regulation of root growth (Acosta et al., 2013; Gasperini et al., 2015), and together with topoisomerase II-associated protein PAT1H1, NINJA participates in the maintenance of root stem cell niche (Yu et al., 2016). In this study, we found a novel function for NINJA in the negative regulation of ERF19 (Figure 8). The repression mechanism(s) of NINJA on ERF19 might be linked to ERF19 association with NINJA that in turn recruits co-repressors such as TPL (Pauwels et al., 2010), and thus suppresses the transcription of ERF19-bound loci. In addition, association with NINJA may change the conformation of ERF19 and subsequently inhibit the transcriptional function of ERF19 as observed in MYC3-JAZ9 regulation (Zhang et al., 2015). Such conformational change may hinder the ability of ERF19 to recruit co-activators and/or to bind to DNA. Our data provide strong evidences that NINJA is involved in the regulation of ERF19 function and further suggest that through modulation of ERF19 at transcriptional and post-translational levels, plants can fine-tune PTI to cope with the vast variety of environmental stimuli they face.

## METHODS

### Biological Materials and Growth Conditions

Growth conditions of *Arabidopsis thaliana* (L. Heyhn.) and *N. benthamiana* were described previously (Yeh et al., 2016). *Arabidopsis* ecotype Col-0 was used as the wild-type for the experiments unless stated otherwise. We obtained mutants *npr1-1* from X. Dong (Duke University, Durham, NC, USA), *ein2-1* from the Arabidopsis Biological Resource Center (https://abrc.osu.edu/), *coi1-16* (Col-6 background) from J.G. Turner (University of East Anglia, Norwich, UK), and *ninja-1* from E.E Farmer (University of Lausanne, Switzerland). *Arabidopsis* transgenic line 35S:GFP was obtained from K. Wu (National Taiwan University, Taipei, Taiwan). The fungus *B. cinerea* was obtained from C.-Y. Chen (National Taiwan University, Taipei, Taiwan) and was grown on potato dextrose broth (PDB)-agar plates in the growth chamber where *Arabidopsis* plants were grown (Zimmerli et al., 2001). Bacterium *Pst* DC3000 was provided by B.N. Kunkel (Washington University, St. Louis, Missouri, USA) and was grown at 28°C, 200 rpm in King’s B medium with 50 mg L^-1^ rifampicin.

### Preparation of Chemicals

Chitin (#C9752, Sigma) was ground to fine powder and suspended in deionized water to make 1 mg mL^−1^ stock solution. The flg22 and elf18 peptides were purchased from Biomatik and dissolved in deionized water to make 1 mM stock solutions. A stock of 10 mM β-Est (#E2758, Sigma) was prepared in DMSO and stored at −20 °C in small aliquots.

### Pathogen Infection Assays

Droplet-inoculation with *B. cinerea*, and assessment of disease symptoms were performed as previously described (Catinot et al., 2015), except 8 μL of *B. cinerea* inoculum per leaf were used in this study. For spray-inoculation with *B. cinerea*, the spore suspension (10^5^ spores mL^-1^ in 1/4 PDB) was evenly sprayed on the leaves of 4-week-old plants until run-off occurred. The infected plants were kept at 100% relative humidity, and disease development was scored at 5 days post inoculation (dpi). Dip-inoculation with *Pst* was performed as previously described (Yeh et al., 2016). To measure bacterial populations, at least 3 plants were used for each experiment. For each plant, five leaf discs (0.2827 cm^2^ /disc) were punched out from 5 different leaves 2 days after *Pst* inoculation. Leaf discs were homogenized in 10 mM MgSO_4_ with mortars and pestles. Three 10-μL droplets of appropriate dilutions were applied on LB plates containing 50 mg L^-1^ of rifampicin. Bacterial colonies were counted after 48 h at 28°C. To assess PTI-mediated resistance to *Pst*, assays were performed as previously described with small modifications (Liu et al., 2015b). Briefly, five leaves per plant were syringe-infiltrated with deionized water or 10 nM elf18. Excessive solution was removed from the leaves with tissue paper. Six hours after treatment, plants were syringe-infiltrated with 10^6^ cfu mL^-1^ *Pst* solution and kept at 100 % relative humidity overnight. Bacterial titers were determined at 2 dpi as mentioned above.

### Generation of Transgenic Plants

The CDS of *ERF19* without stop codon was amplified from Col-0 cDNA with ERF19-F1 and ERF19-R1 primers and cloned into pCR8-TOPO vector (Invitrogen) to create pCR8-ERF19. After sequence confirmation, *ERF19* CDS was sub-cloned into pMDC83 (Curtis and Grossniklaus, 2003) and pEarleyGate103 (Earley et al., 2006) vectors via LR reaction to create pMDC83-ERF19 and pEarleyGate103-ERF19 constructs. To create the inducible construct, ERF19-GFP CDS was partially digested from pEarleyGate103-ERF19 with XhoI and PacI. The ERF19-GFP fragment was ligated with pMDC7 vector (Zuo et al., 2000; Curtis and Grossniklaus, 2003) and digested with XhoI and PacI to create pMDC7-ERF19. To construct chimeric ERF19-SRDX, the genomic fragment of *ERF19* including its promoter region (−1 to −1535 bp) was amplified by PCR using ERF19-F2 and ERF19-R2 primers. The product was digested with Asc1 and Sma1 and then introduced into the same enzyme-treated VB0227 vector. After confirming inserted region sequences, completed *ProERF19:ERF19-SRDX:HSP* part was transferred into pBCKH(VB0047) (Mitsuda et al., 2011) binary vector by LR reaction to create pBCKH-ERF19-SRDX. *Agrobacterium tumefaciens* GV3101 was used to deliver the constructs into plants (Martinez-Trujillo et al., 2004). Constructs pMDC83-ERF19, pEarleyGate103-ERF19, pMDC7-ERF19, and pBCKH-ERF19-SRDX were used to generate transgenic ERF19-OE, ERF19-OE/*ninja-1*, ERF19-iOE, and ERF19-SRDX lines, respectively. Independent homozygous T3 lines with a single T-DNA insertion were used for the experiments. Primers used to construct the plasmids are summarized in Supplemental Table 1.

### Treatment of β-Est

To examine *ERF19* or ERF19-GFP expression induced by β-Est, 12-day-old seedlings were submerged in liquid 1/2 MS supplemented with 20 μM β-Est or DMSO. After 24 h of incubation, seedlings were collected in liquid nitrogen for downstream analyses. To evaluate resistance to *B. cinerea*, 20 μM β-Est or DMSO was syringe-infiltrated into the leaves of 5-week-old plants, and the treated plants were placed back to their normal growth conditions. After 24 h, the plants were droplet-inoculated with *B. cinerea* spores as mentioned above.

### Subcellular Localization

Twelve-day-old ERF19-iOE1 and 35S:GFP transgenic lines were incubated in liquid 1/2 MS supplemented with 20 μM β-Est. After 24 h incubation, the seedlings were vacuum infiltrated with DAPI solution (5 μg mL^-1^) for 2 min and washed 3 times with distilled water. The GFP and DAPI signals in the roots were imaged with Zeiss LSM 780 confocal microscope.

### RT-PCR

To monitor MAMP-or pathogen-induced *ERF19*, 12-day-old seedlings were incubated in liquid 1/2 MS one night before treatments with 200 μg mL^-1^ chitin, 100 nM flg22, 100 nM elf18, 5 × 10^5^ *B. cinerea* spores mL^-1^, or 10^7^ cfu mL^-1^ *Pst*. To examine PTI marker genes, 12-day-old seedlings incubated in 1/2 MS overnight were treated with 200 μg mL^-1^ chitin, 1 μM flg22, or 1 μM elf18. Samples were collected in liquid nitrogen at the indicated time points. Total RNA isolation, reverse transcription, and real-time PCR analyses were performed as described (Catinot et al., 2015). The gene *UBIQUITIN 10* (*UBQ10*, AT4G05320) was used for normalization. For RT-PCR, two microliters of cDNA was used as template, and standard PCR conditions were applied as described (Huang et al., 2014). *UBQ10* was used as a loading control. Primers used are listed in Supplemental Table 1.

### Callose Deposition Assays

Fourteen-day-old seedlings were incubated in liquid 1/2 MS one night before treatments with 200 μg mL^-1^ chitin, 100 nM flg22, 100 nM elf18, or deionized water. Twenty-four hours later, the liquid media were replaced with 95 % ethanol to clear chlorophyll. The cleared seedlings were washed twice with 70 % ethanol and then 3 times with distilled water. The seedlings were then vacuum-infiltrated with 0.1 % aniline blue in 150 mM K_2_ HPO_4_ (pH 10.5/KOH) and incubated in the dark for 2 h. Callose deposits in the cotyledons were visualized under UV illumination using a Nikon Optiphot-2 microscope. Quantification of callose spots was performed as described (Kohari et al., 2016).

### Protoplast Preparation and Transfection

*Arabidopsis* protoplasts were prepared from the leaves of 5-week-old plants as previously described (Wu et al., 2009). Polyethylene glycol-mediated protoplast transfection was performed as described (Yoo et al., 2007).

### PTA Assays

Protoplast transactivation assays were performed as previously described (Hsieh et al., 2013). A plasmid carrying the gene encoding firefly luciferase (fLUC) under the control of UAS targeted by the yeast GAL4 TF were used as the reporter plasmid. The reference plasmid carries the gene encoding Renilla luciferase (rLUC) under the control of 35S promoter. Effector plasmid harboring the DNA binding domain of GAL4 expressed from the 35S promoter was used as the empty vector control (GAL4DB). The fragment of *ERF19* CDS amplified by PCR using ERF19-F3 and ERF19-R3 primers was digested with XmaI and SalI. The digested fragment was then introduced into GAL4DB digested with the same enzymes to create GAL4DB-ERF19 effector plasmid. To construct GAL4DB-ERF19-SRDX effector plasmid, the fragment ERF19-SRDX, amplified from pBCKH-ERF19-SRDX with primers ERF19-F and SRDX-R, was ligated with the vector backbone, amplified from GAL4DB with primers pGAL4-F and pGAL4-R, by blunt-end cloning. The CDS of *NINJA* was amplified from Col-0 cDNA with NINJA-F and NINJA-R1 primers and introduced into pCR8-TOPO vector. The insert was then sub-cloned into pGWHA, a plasmid modified from p2FGW7 (Karimi et al., 2002), by substituting the GFP tag with a single HA tag, by LR reaction to create the NINJA effector plasmid. One hundred microliters of 2 × 10^4^ protoplasts mL^-1^ were transfected with 5 μg, 4 μg, and 1 μg of effector plasmids, reporter plasmids, and reference plasmids, respectively. After 24 h, the transfected protoplasts were harvested and the luciferase activities were analyzed using the Dual-Luciferase Reporter Assay System (Promega). Data are presented as the normalized fLUC activities relative to the no effector control (set as 1). Primers used to construct the plasmids are summarized in Supplemental Table 1.

### MAMP-Induced Growth Inhibition

Growth inhibition experiments were performed as described (Ranf et al., 2011). Briefly, ten 5-day-old seedlings of the same genotype were transferred into 6-well-plates supplemented with liquid 1/2 MS (0.5 g L^-1^ MES, 0.25 % sucrose, pH 5.7). The seedlings were treated with water or MAMPs at indicated concentration. The treated seedlings were further grown for another 10 days under normal growth conditions. Ten seedlings in a single well were blotted dry on tissue paper and weighed as a whole.

### Co-IP Assay in *Arabidopsis* Protoplasts

Full-length CDS of *NINJA* without stop codon, amplified with NINJA-F and NINJA-R2 primers (Supplemental Table 1), was cloned into pCR8-TOPO entry vector and sub-cloned into pEarleyGate103 destination vector with 35S promoter and C-terminal GFP fusion (Earley et al., 2006) by LR reaction. The GFP empty vector control was created by digesting pEarleyGate103 with XhoI to remove the Gateway cassette. The *ERF19* CDS from pCR8-ERF19 was first introduced into pGWB14 vector with C-terminal triple HA fusion (Nakagawa et al., 2007) via LR reaction, and the fragment 35S:ERF19-HA_3_:NOS was amplified with 35S-F and NOS-R primers by PCR (Supplemental Table 1). This fragment was cloned into pCR8-TOPO to create a plant expression plasmid with high copy number. Protoplast transfection and Co-IP were performed as previously described (Yeh et al., 2015).

### BiFC in *N. benthamiana*

Using LR reaction, *ERF19* CDS without stop codon and *NINJA* CDS with stop codon were introduced into pVYNE with C-terminal fusion of YFP N-terminus (nYFP) and pVYCE(R) with N-terminal fusion of YFP C-terminus (cYFP) (Waadt et al., 2008), respectively. To create the construct for the nuclear marker, the nuclear localization signal (NLS) was fused to the N-terminus of mCherry by PCR with primers NLS-mCherry-F and mCherry-R (Supplemental Table 1). This fragment was cloned into pENTR/D-TOPO vector and digested with SmaI to create a blunt-end vector. The vector was then ligated with the PCR fragment mCherry-NLS, amplified with primers mCherry-F and mCherry-NLS-R (Supplemental Table 1), to create the complete NLS-mCherry-mCherry-NLS sequence. This sequence was introduced into pEarleyGate100 (Earley et al., 2006) by LR reaction to create the construct for the nuclear marker. The constructs were transformed into *A. tumefaciens* GV3101 by electroporation. For transient expression, *Agrobacteria* were grown in LB medium (28 °C, 200 rpm) supplemented with appropriate antibiotics overnight. Cells were pelleted and resuspended in 10 mM MgCl_2_, 10 mM MES, and 200 μM acetosyringone. After 3 h of incubation on bench top, *Agrobacterium* strains carrying the BiFC constructs were mixed 1:1 to a final OD_600_ of 0.4 for each strain, and the nuclear marker strain was added to a final OD_600_ of 0.1. The inocula were syringe infiltrated into 3-week-old *N. benthamiana* leaves. Two days later, the transformed leaves were imaged with Zeiss LSM 780 confocal microscope.

### Protein Extraction in *Arabidopsis* Seedlings

Extraction of total proteins from *Arabidopsis* seedlings was performed as previously described (Tsugama et al., 2011). Briefly, 10-day-old seedlings were weighed and homogenized in lysis buffer [0.1 M EDTA (pH 8.0), 0.12 M Tris-HCl (pH 6.8), 4 % SDS, 10 % beta-mercaptoethanol, 5 % glycerol, and 0.005 % bromophenol blue] at a ratio of 1 mg fresh weight per 10 μL lysis buffer. Homogenized samples were boiled for 10 min and centrifuged at 12,000 g for 5 min. The supernatants were directly used for immunoblotting.

### SDS-PAGE and Immunoblotting

For immunoblotting, protein samples were resolved on 10 to 12 % SDS-PAGE gels run at 80-140 V for 2 h. Resolved proteins were transferred to PVDF membranes (Millipore) at 100 V for 1 h at 4 °C. Membranes were blocked by 5 % nonfat milk in TBST for 2 h. Blocked membranes were incubated with primary antibodies anti-GFP (#sc-9996, Santa Cruz Biotechnology) or anti-HA (#sc-7392, Santa Cruz Biotechnology) overnight. Membranes were washed 3 times in TBST before 1 h incubation with secondary antibodies anti-mouse-HRP (#sc-2005, Santa Cruz Biotechnology) or anti-rabbit-HRP (#sc-2004, Santa Cruz Biotechnology). Signals were detected by an enhanced chemiluminescence system (WesternBright Sirius HRP substrate, Advansta) and a scanner (Wealtec Corp.) following manufacturers’ instructions.

### Y2H Assays

Full length CDS of *ERF19* and *NINJA* were sub-cloned into gateway vectors pGADT7 and pGBKT7, respectively via the LR reaction. The constructs were transformed into yeast strain AH109 based on the LiAc-mediated transformation protocol following the manufacturer’s instructions (Clonetech). At least 10 co-transformed yeast colonies were plated on Synthetic Drop-Out (SD) medium supplemented with X-α-Gal (Clonetech) but without leucine, tryptophan, and histidine (-L-W-H). The plates were incubated at 30 °C for 3 days to test the nutritional marker gene expression and galactosidase activity of the MEL1 reporter protein.

## Accession Numbers

Sequence data from this article can be found in the *Arabidopsis* Genome Initiative under the accession numbers: *ERF19* (AT1G22810), *NINJA* (AT4G28910), *UBQ10* (AT4G05320), *PDF1.2* (AT5G44420), *PDF1.3* (AT2G26010), *PR1* (AT2G14610), *PR2* (AT3G57260).

## Supplemental Data

**Figure 1.** Characterization of the HA-ERF19 line.

**Figure 2.** Characterization of lines overexpressing *ERF19*

**Figure 3.** *B. cinerea*-mediated lesions in ERF19-iOE lines.

**Figure 4.** Growth phenotypes of ERF19-OE and ERF19-iOE lines.

**Figure 5.** *ERF19* up-regulation by chitin in defense signaling mutants.

**Figure 6.** Expression of PTI marker genes in ERF19-OEs.

**Figure 7.** Characterization of ERF19-SRDXs.

**Figure 8.** Characterization of ERF19-OEs/*ninja-1*.

**Table 1.** Primers used in this study.

## ACKNOWLEDGEMENTS

We thank ABRC, X. Dong, J.G. Turner, E.E Farmer, and K. Wu for providing seeds, and C.-Y. Chen and B.N. Kunkel for *B. cinerea* and *Pst* bacteria, respectively. We also acknowledge members of Zimmerli laboratory for critical comments. We thank the Technology Commons (TechComm), College of Life Science, National Taiwan University for providing qRT-PCR equipment and excellent technical assistance with the confocal laser scanning microscopy. This work was supported by the Ministry of Science and Technology of Taiwan, grants 98-2311-B-002-008-MY3, 102-2311-B-002-027, 103-2311-B-002-004, and 104-2311-B-002-003 (to L.Z.).

## AUTHOR CONTRIBUTIONS

P.-Y. and L.Z. designed the research. P.-Y. performed most experiments. Y.-P. L. started the project by screening the *At*TORF-Ex collection. J.C. generated ERF19-iOE lines and performed Co-IP analyses. B. J. conducted Y2H and BiFC experiments and rough phenotyping of ERF19-OE/*ninja-1* lines. J.-H Y. performed flg22-induced *ERF19* analysis. K.C. cloned ERF19-SRDX and generated ERF19-SRDX lines. P.-Y. and L.Z. analyzed the data and wrote the manuscript. L.Z. supervised the project.

